# Genomic regions associated with chocolate spot (*Botrytis fabae* Sard.) resistance in faba bean (*Vicia faba* L.)

**DOI:** 10.1101/2021.11.22.469473

**Authors:** Tadesse S. Gela, Margaret Bruce, Wei Chang, Frederick L. Stoddard, Alan H. Schulman, Albert Vandenberg, Hamid Khazaei

## Abstract

Chocolate spot (CS), caused by *Botrytis fabae* Sard., is an important threat to global faba bean production. Growing resistant faba bean cultivars is, therefore, paramount to preventing yield loss. To date, there have been no reported quantitative trait loci (QTLs) associated with CS resistance in faba bean. The objective of this study was to identify genomic regions associated with CS resistance using a recombinant inbred line (RIL) population derived from resistant accession ILB 938. A total of 165 RILs from the cross Mélodie/2 × ILB 938/2 were genotyped and evaluated for CS reactions under replicated controlled climate conditions. QTL analysis identified five loci contributing to CS resistance on faba bean chromosomes 1 and 6, accounting for 5.0–23.4% of the total phenotypic variance. The sequences of SNP markers linked to resistance QTLs on chromosome 1 that have the largest effects encode multiple classes of putative disease and/or defense-related genes. The results of this study not only provide insight into disease-resistance QTLs, but also can be used as potential targets for marker-assisted breeding in faba bean genetic improvement for CS resistance.

**Key message:** QTL mapping identified key genomic regions associated with chocolate spot resistance on faba bean chromosomes 1 and 6, which may serve as novel genetic tools to develop disease-resistant faba bean cultivars.

## Introduction

Faba bean (*Vicia faba* L., 2n=2x=12) is a widely grown diploid cool-season legume (pulse) crop with a genome size of approximately 13 Gb (Soltis et al. 2003). It is the fifth most important pulse crop, after common bean (*Phaseolus vulgaris* L.), chickpea (*Cicer arietinum* L.), pea (*Pisum sativum* L.), and lentil (*Lens culinaris* Medik.), with an annual production of 5.4 Mt (FAOSTAT 2021). It provides an affordable source of dietary protein, fiber, carbohydrates, and other valuable nutrients (Duc et al. 2015; Khazaei and Vandenberg 2020). The faba bean seed contains about 30% protein (Warsame et al. 2018), making it one of the best alternative sources for the plant-based protein industry for food, feed, and extractable protein. The crop plays an important role in a sustainable agriculture system, due to its symbiotic fixation of atmospheric nitrogen and its use as a break crop in cereal-dominated rotations (Angus et al. 2015). Global faba bean production is generally challenged by foliar diseases such as ascochyta blight (*Ascochyta fabae* Speg.), chocolate spot (*Botrytis fabae* Sard.), rust (*Uromyces viciae fabae* Pers.), downy mildew (*Peronospora viciae* (Berk.) Caspary), and gall disease (*Physioderma viciae*) (Adhikari et al. 2021).

Chocolate spot (CS), caused by the necrotrophic fungal pathogen *Botrytis fabae* (Sard.), is one of the most destructive diseases of faba bean plants worldwide and can cause up to 90% yield reduction on susceptible cultivars when conducive environmental conditions prevail (Gorfu and Yaynu 2001; Bouhassan et al. 2004; Tivoli et al. 2006; Beyene et al. 2018). CS can manifest itself on the above-ground parts of the plant at any growth stage, mainly on leaves. It initially appears on leaves as reddish-brown spots that develop into chocolate-colored lesions. These can enlarge into extensive necrotic zones that lead to severe premature defoliation. If the disease develops at flowering time, it can cause complete crop failure. The pathogen survives on crop residues or lodged stubble of the previous year’s growing season and on contaminated seeds (Harrison 1988). Its spores are spread primarily between fields by wind and rain. Variation in virulence among *B. fabae* isolates has been demonstrated (Hutson and Mansfield 1980), although no race classification has been described so far. The strategies used to control chocolate spot on faba bean include crop rotation, reduced planting density, timely fungicide applications, use of clean seeds, and host-plant resistance (Stoddard et al. 2010). Other species of *Botrytis* contribute to chocolate spot disease, but *B. fabae* is the main one (Fan et al. 2015).

Genetic resistance is a key part of any integrated disease management approach to prevent yield loss caused by CS. Fungal evolution and migration leads to the need for continuous incorporation of new sources of resistance genes or alleles into elite breeding materials. A few sources of partial genetic resistance to CS have been identified in faba bean germplasm (Hanounik and Robertson 1988; Bouhassan et al. 2004; Villegas-Fernández et al. 2009; Maalouf et al. 2016). Partial resistance to CS was transferred into elite breeding lines and resulted in the release of several cultivars with moderate levels of resistance (Temesgen et al. 2015; Maalouf et al. 2019). Incorporation and pyramiding of CS resistance genes from multiple sources into elite cultivars could be facilitated with marker-assisted selection (MAS). However, the use of MAS in faba bean CS resistance breeding is limited by lack of knowledge of the genetic basis of the resistance so-far identified, by the presence of pathogen variability in the major faba bean diseases (Hanounik and Robertson 1988; Gorfu 1996), and by the lack of adequate genomic resources.

In recent years, progress has been made in developing single nucleotide polymorphism (SNP)-based genetic maps of faba bean (e.g. Ellwood et al. 2008; Cruz-Izquierdo et al. 2012; Kaur et al. 2014; Webb et al. 2016; Carrillo-Perdomo et al. 2020). Among these maps, the densest SNP-based consensus maps are reported by Webb et al. (2016), constructed from six mapping populations using 687 SNP markers, and Carrillo-Perdomo et al. (2020), a consensus map of three mapping populations consisting of 1,728 SNP markers distributed in six linkage groups. Concurrently, several QTL studies for various faba bean improvement traits have been carried out using different genetic maps (see reviews by O’Sullivan and Angra 2016; Maalouf et al. 2019; Khazaei et al. 2021a) for major foliar diseases such as ascochyta blight (e.g. Avila et al. 2004; Kaur et al. 2014; Atienza et al. 2016; Sudheesh et al. 2019) and rust (e.g. Adhikari et al. 2016; Ijaz et al. 2021). Despite the importance and widespread global nature of CS, no QTL studies have been published to our knowledge. In addition, breeding for CS resistance has been difficult because screening for the disease under field conditions is unreliable, especially in dry environments where faba bean is grown mostly as a rain-fed crop (Adhikari et al. 2021). Humid conditions, which are uncommon in many faba bean-growing areas, provide the opportunity to screen in the field. Identification of reliable molecular markers for use in MAS systems for improving CS resistance is essential for future faba bean breeding. The objective of this study was to identify QTLs associated with CS resistance using an advanced RIL mapping population developed from accession ILB 938 which has partial resistance to CS (Khazaei et al. 2018) under climate-controlled conditions.

## Material and methods

### Plant material

Genetic resistance to *B. fabae* was evaluated using 165 RILs from the Mélodie/2 × ILB 938/2 cross (Khazaei et al. 2014a). The RILs were advanced using single seed descent to the F_8_ generation. Then the F_8_-derived bulked seeds of the RILs were selfed for at least two additional generations before disease phenotyping. Mélodie/2 was bred at INRA (France) as a source of low vicine-convicine (Khazaei et al. 2014a). ILB 938 (IG 12132) originates from the Andean region of Colombia and Ecuador and has resistance to several biotic and abiotic stresses, including CS (Maalouf et al. 2016; Khazaei et al. 2018). It has high water use efficiency (Link et al. 1999; Khazaei et al. 2014a) and carries a gene that decouples pigmentation in flowers from that in stipules (Khazaei et al. 2014b). It was the first identified source of resistance for CS in faba bean (Hanounik 1982; Khalil et al. 1984; Robertson 1984). ILB 938 resulted from mass selection within ILB 438 (IG 11632). ICARDA’s registered BPL (pure line) derivatives of ILB 438 and ILB 938 are BPL 710 and BPL 1179, respectively.

### Inoculation and phenotyping for chocolate spot reactions

A culture stock of *B. fabae* isolate FB-7 was obtained from the pulse pathology laboratory, University of Saskatchewan for mass spore production. Conidia were cultured on faba bean extract media plates (18 g of agar, 20 g of dextrose, 20 g of NaCl, and 1 L of faba bean extract which is prepared by boiling 400 g of faba bean seeds in 1 L of distilled water for 45 minutes and diluting with 1 L of distilled water after removing the faba bean seeds) and incubated for 14 d at room temperature in a cycle of 12 h of darkness and 12 h of light to induce sporulation. Plates were then flooded with sterile deionized water and conidia were harvested by scraping the colonies with the edge of a sterile glass microscope slide. The suspension was collected and filtered through one layer of cheesecloth into a clean Erlenmeyer flask. The concentration of the conidial suspension was adjusted to 5 × 10^5^ spores mL^-1^ using a haemocytometer. The surfactant Tween 20 (polyoxyethylene sorbitan monolaurate) was added at the rate of 1 to 2 drops per 1000 mL of suspension, and the suspension was shaken well before inoculation.

The parents and 165 RILs were grown in a climate-controlled growth chamber (GR48, Conviron, Winnipeg, Canada) at the University of Saskatchewan’s College of Agriculture and Bioresources phytotron facility, Saskatoon, Canada. The growth chamber conditions were adjusted to 18 h light and 6 h dark, with the temperatures maintained at 21 °C (day) and 18 °C (night), and the photosynthetic photon flux density was set to 300 μmol m^-2^ s^-1^ during the light period at the crop canopy level. Three seeds of each genotype were sown in individual 3.8 L pots (15.5 cm in diameter) containing a soil-less mixture (Sunshine Mix No. 3, Sun Grow Horticulture® Ltd., Vancouver, BC, Canada) and fertilized once per week using 3 g L^-1^ of soluble N:P:K (20:20:20) PlantProd® fertilizer (Nu-Gro Inc., Brantford, ON, Canada). Three weeks after seeding, the plants were inoculated with a spore suspension (5 × 10^5^ spores mL^-1^) at ∼3 mL per plant until runoff using a pressurized knapsack sprayer. Plants were placed in an incubation chamber for seven days. Two humidifiers (Vicks Fabrique Paz Canada, Inc., Milton, ON, Canada) were placed in the incubation chamber to ensure 90–100% relative humidity for infection and disease development. The disease severity data were collected per individual plant at seven days post-inoculation (dpi) using a 0 to 10 rating scale with 10% increments. The experiment was repeated four times under the same growing conditions during 2020-2021, and each time disease scores were recorded from three replicates (one plant per replicate). Data were converted to percentage disease severity using the class midpoints for data analysis and analyzed as a randomized complete block design.

### Statistical analysis

Disease score data were analyzed using SAS v.9.4 (SAS Institute, Cary, USA). Normality and variance homogeneity of the residuals were analysed using the Shapiro-Wilk normality test and Levene’s test for homogeneity, respectively. Genotypes were treated as fixed effects and blocks as random effects; significance of variances were declared at 5% significance level. Least square means were estimated for genotype using LSMEANS statements and used for QTL analysis. Multivariate relationships between RILs and experimental repeats were investigated by principal component analysis (PCA) and biplots of autoscaled (i.e., each repeat mean and grand mean centred at the origin) data in R (R Core Team 2020) with the ‘factoextra’ package.

### Genotyping

Genomic DNA was extracted from three-day-old germinated embryo axes of the parents and 188 RILs from the mapping population using the CTAB method, as described previously (Björnsdotter et al. 2021). DNA quality was assessed by agarose gel electrophoresis; concentrations were determined with a Quant-iT PicoGreen dsDNA Assay Kit (ThermoFisher Scientific, UK) following the manufacturer’s guidelines. DNA samples were brought to a concentration of 35 ng/µl in 40 µl aliquots and genotyped using the Axiom ‘Vfaba_v2’ 60K array that was developed from metatranscriptome data (O’Sullivan et al. 2019).

### Linkage map construction

Genotypic data for the RIL population was filtered against markers showing significant segregation distortion (deviating from the expected 1:1 ratio) using a chi-square (*χ*^*2*^) test. Markers missing in 1% or more of the data were removed from the analysis. Draft maps were generated from the remaining 180 RILs using ASMap software (Taylor et al. 2017) with a p-value of 1E^-10^ and a maximum distance between SNP markers of 15.0 cM for grouping them into linkage groups. This linkage map was refined using MapDisto v. 1.7.7.0.1 (Lorieux 2012) with a logarithm of odds (LOD) score of 3.0 and a cut-off recombination value of 0.35. The best marker order was estimated using both “AutoCheckInversions” and “AutoRipple” commands in MapDisto. Distances between markers were calculated using the Kosambi function (Kosambi 1943). Linkage groups (LGs) were scanned and corrected for double recombinants using MapDisto v. 1.7.7.0.1 (Lorieux 2012). The final LGs were assigned to chromosomes according to the NV644 × NV153 genetic map developed at the University of Reading, UK (unpublished).

### QTL mapping of disease resistance

Windows QTL Cartographer 2.5 software (Wang et al. 2012) and multiple QTL mapping (MQM, Manichaikul et al. 2009), run in R/qtl software (Broman et al. 2003), were used to detect QTLs. For Windows QTL Cartographer, composite interval mapping (CIM) was implemented using the Kosambi map function with Ri1 cross type (recombinant inbred line, derived by selfing) at a 1.0 cM interval walk speed. Cofactor selection was performed using forward and backward regression in the standard CIM model with a probability of in and out of 0.1 and a window size of 5 cM. QTL significance thresholds were determined by permutation tests (1000 permutations) at a significance level of *P* = 0.05. Multiple QTL mapping was completed with the “*stepwiseqtl*” function (Broman et al. 2003), using the Haley-Knott regression (Haley and Knott 1992) methods. The optimal QTL model was chosen based on the highest penalized LOD score (Manichaikul et al. 2009) after forward and the backward selection and elimination modeling. Penalties for model selection and the genome-wide significance threshold (= 0.05) were determined through 1000 permutations of the “*scantwo*” function for two-dimensional QTL scanning. The percentage of the phenotypic variance explained (PVE) and effects of QTLs were obtained by fitting a mixed linear model using the “*fitqtl*” function. The confidence intervals for each QTL were estimated using the “*lodint*” function that calculates the 1.5 LOD support intervals. To identify putative candidate genes, the sequences of the SNP markers in the QTL intervals were searched by BLASTn in Phytozome v13 (Goodstein et al. 2012) on the *Medicago truncatula* reference genome (Mt4.0v1). QTL positions on the linkage map were drawn by MapChart v. 2.2 (Voorrips 2002).

## Results

### Phenotypic analysis of the parental lines and RIL population

Individuals in the RIL population showed significant variation in CS resistance (F-value = 2.21, *P* = 0.0001). The parental lines Mélodie/2 and ILB 938/2 exhibited significant differences in resistance to CS under growth chamber conditions (Fig 1). ILB 938/2 had a partial resistant reaction with mean disease severity of 30.4%, while Mélodie/2 showed a moderately susceptible reaction with a mean of 53.3%. The distribution of disease severity as a measure of CS response for RILs showed continuous variation ranging from 20.8% to 62.9%, with a mean of 44.5% (Fig 1), suggesting polygenic regulation of CS severity. Similarly, the autoscaled PCA indicated that the data had no underlying structure for the first two principal components (PC1 vs PC2), which explained approximately 66.1% of the variance. The PCA-biplot graph also displayed a strong positive correlation among the four screening repeats, which were represented well by the graph (i.e., the vector with equal length), demonstrating that the data could be used for accurate mapping of CS resistance QTLs (Fig 2).

**Figure 1.**
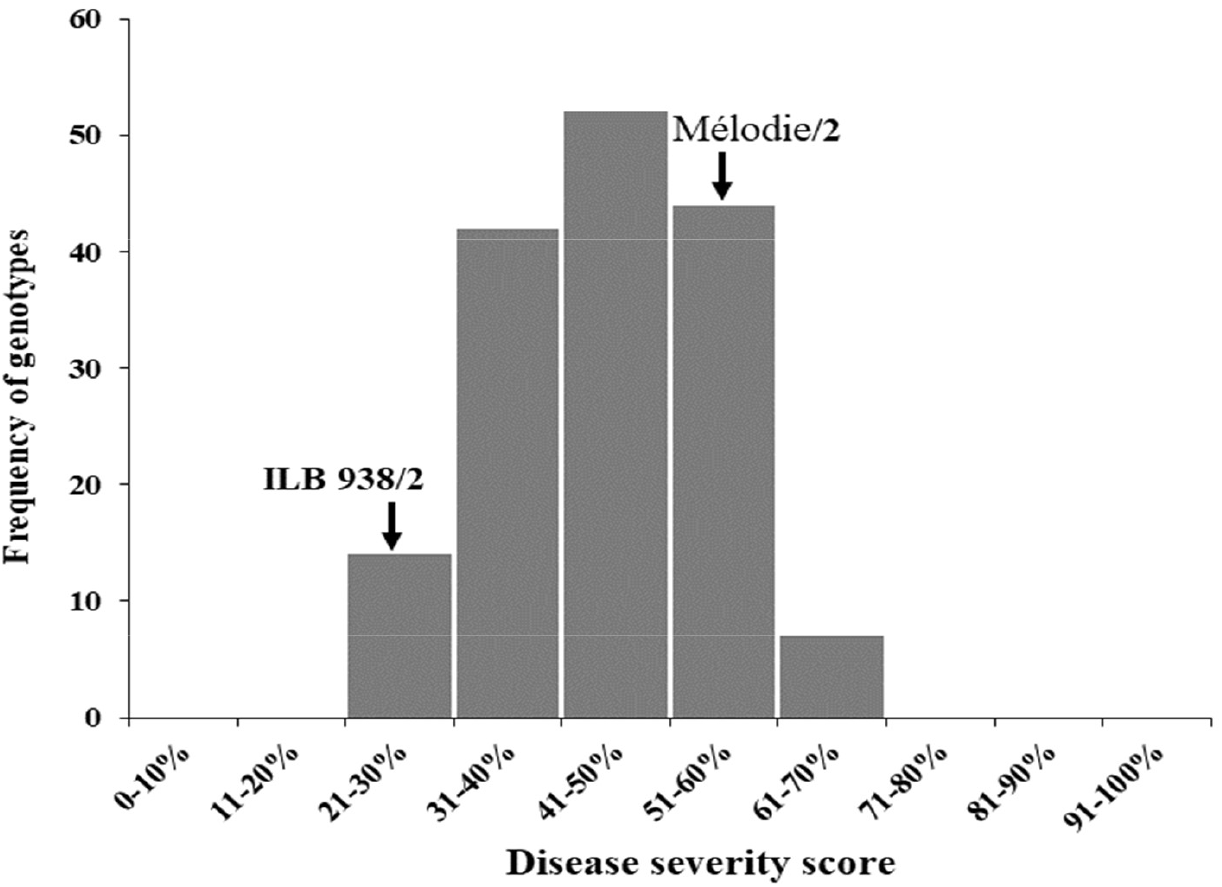
Frequency distributions of chocolate spot disease scores in 165 RILs derived from cross Mélodie/2 × ILB 938/2 evaluated under growth chamber conditions. Each value is the mean of 4 screenings, each with 3 replicates. The arrows indicate the average disease severity of the parents. Disease severity was rated on a 0–10 scale, where the disease severity score increased in 10% increments

**Figure 2.**
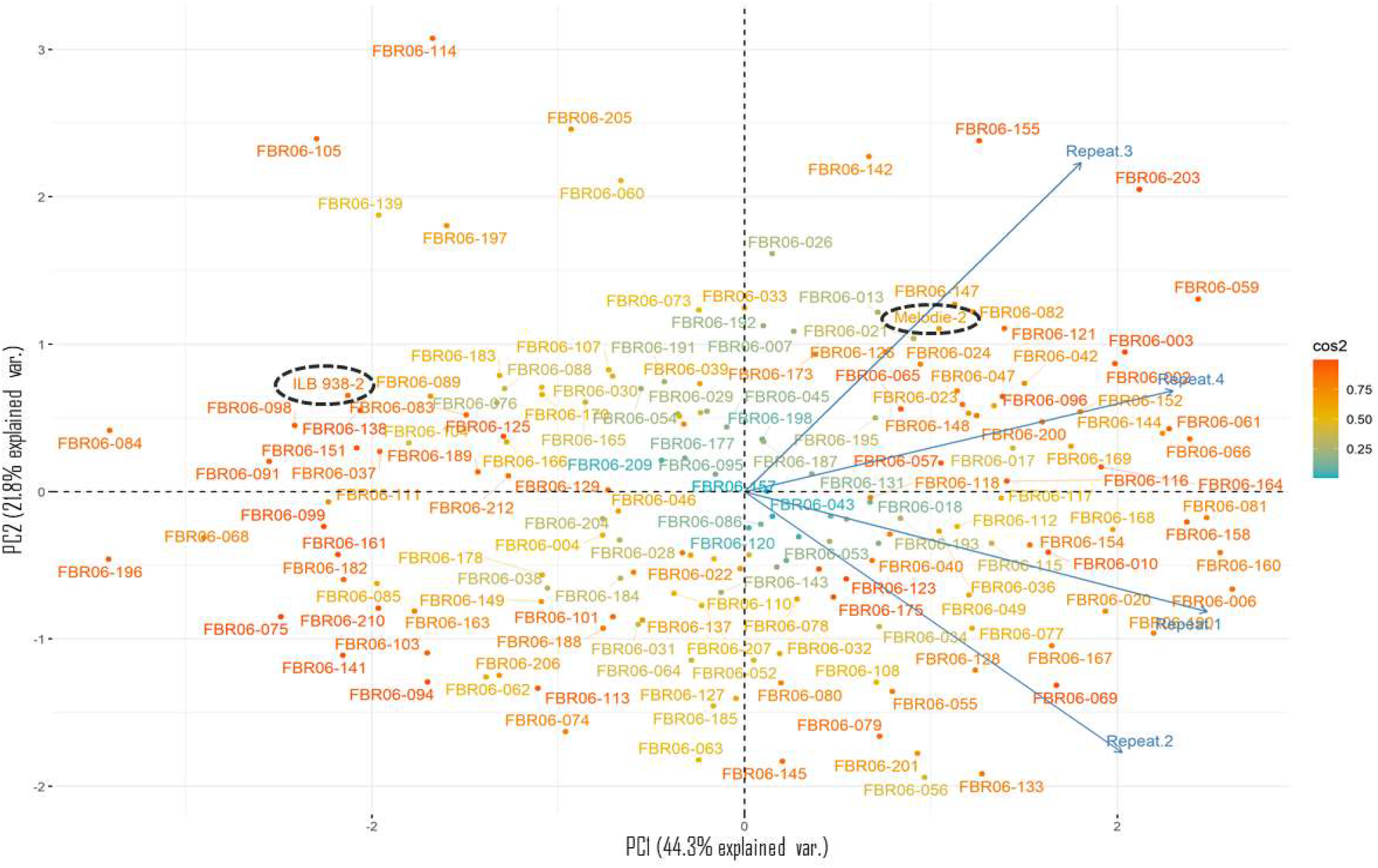
PCA-biplot graph of PC1 vs PC2 for auto-scaled and centered data for chocolate spot severity for each repeat (rep) collected from a 165 recombinant inbred lines (RILs) population of a cross Mélodie/2 × ILB 938/2 cross (FBR-06). Each RIL score, which is a deviation from its rep average value, was displayed as a point on the graph and reps as a blue arrow (vector). The parental lines are circled with black dots. The origin of the graph represents the average value for both each rep and across all repeats. The angle between any two arrows indicates the correlation between repeats (if the angle is 90 degrees, it means both variables show a lack of correlation) and the length of the arrow represents the variance explained by two components (all equal length means a perfect fit). The colour bar indicator on the right shows the deviation of each score from the averages

### Linkage map construction

A total of 35,363 SNP markers were filtered for polymorphism between the parents, significant segregation distortion, and missing data. A final genetic map was constructed from 4,089 SNP markers, which mapped to six linkage groups (LGs) representing the six chromosomes of faba bean (Table 1). The LGs were numbered to match the respective faba bean consensus map based on where the markers lie (Webb et al. 2016). The linkage map spanned a total genetic distance of 1229.5 cM with an average marker interval of 0.3 cM. The genetic distance within LGs varied from 124.2 cM for LG 4 to 396.1 cM for LG 1. LG 4 contained the fewest SNP markers, whereas LG 1 contained the most SNP markers (Table 1; Fig S1).

**Table 1.** Characteristics of the genetic linkage map for the 165 faba bean RIL population derived from cross Mélodie/2 × ILB 938/2 at F8.

| Chromosomes<br>(Linkage groups) | Number of<br>markers | Map length (cM) | Average marker<br>interval (cM) | Maximum gap<br>(cM) |
| --- | --- | --- | --- | --- |
| 1 | 1262 | 396.1 | 0.31 | 3.23 |
| 2 | 656 | 218.1 | 0.33 | 3.21 |
| 3 | 668 | 181.4 | 0.27 | 3.58 |
| 4 | 488 | 124.2 | 0.25 | 2.22 |
| 5 | 488 | 139.8 | 0.29 | 4.32 |
| 6 | 527 | 169.9 | 0.32 | 3.55 |
| Total | 4089 | 1229.5 | 0.30 | - |

### QTL analysis

Five QTLs associated with chocolate spot resistance were detected in the RIL population (Table 2 and Figure 3). Among these, two QTLs (qBF1.1 and qBF1.2) were identified on LG 1 by both QTL mapping methods, CIM and MQM, implemented in WinQTLCart and R/qtl software, respectively. The QTL qBF1.1 accounted for 8.9–10.7% of PVE, whereas qBF1.2 explained 12.2–12.8% of the PVE (Table 2). These QTLs were derived from the resistant line ILB 938/2. On the other hand, the remaining three QTLs, one on LG 1 (qBF1.3), explained 7.3% of the PVE, and two on LG 6 (qBF6.1 and qBF6.2), accounted for 6.2 and 6.3 % of the PVE, respectively, were detected solely by CIM methods. The resistance allele for qBF6.1 was contributed by ILB 938/2, whereas the moderately susceptible parent Mélodie/2 was the donor of qBF1.3 and qBF6.2 (Table 2).

**Table 2.** Quantitative trait loci (QTL) for resistance to chocolate spot in 165 RIL population derived from cross Mélodie/2 × ILB 938/2.

| QTL | LG<sup>a</sup> | Peak<br>(cM) | 2- LOD<br>interval (cM) | CIM<sup>d</sup> |  |  | MQM<sup>d</sup> |  |  |
| --- | --- | --- | --- | --- | --- | --- | --- | --- | --- |
|  |  |  |  | LOD | R<sup>b</sup> (%) | Add<sup>c</sup> | LOD | R<sup>b</sup> (%) | Add<sup>c</sup> |
| qBF1.1 | 1 | 28.5 | 22.7 – 29.7 | 4.2 | 8.9 | -3.0 | 4.5 | 10.7 | -3.2 |
| qBF1.2 | 1 | 127.3 | 125.2 – 130.4 | 5.7 | 12.2 | -3.5 | 5.3 | 12.8 | -3.6 |
| qBF1.3 | 1 | 191.0 | 189.4 – 192.9 | 3.5 | 7.3 | 2.8 | - | - | - |
| qFB6.1 | 6 | 109.5 | 104.1 – 113.2 | 3.7 | 6.2 | -2.5 | - | - | - |
| qFB6.2 | 6 | 164.0 | 159.9 – 168.8 | 3.1 | 6.3 | 4.1 | - | - | - |

<sup>a</sup>Linkage group, <sup>b</sup>R<sup>2</sup> - Percentage of phenotypic variance explained by QTL, <sup>c</sup>Additive genetic effect, <sup>d</sup>CIM - composite interval mapping in WinQTLCart and MQM - multiple QTL mapping in R/qtl software

**Figure 3.**
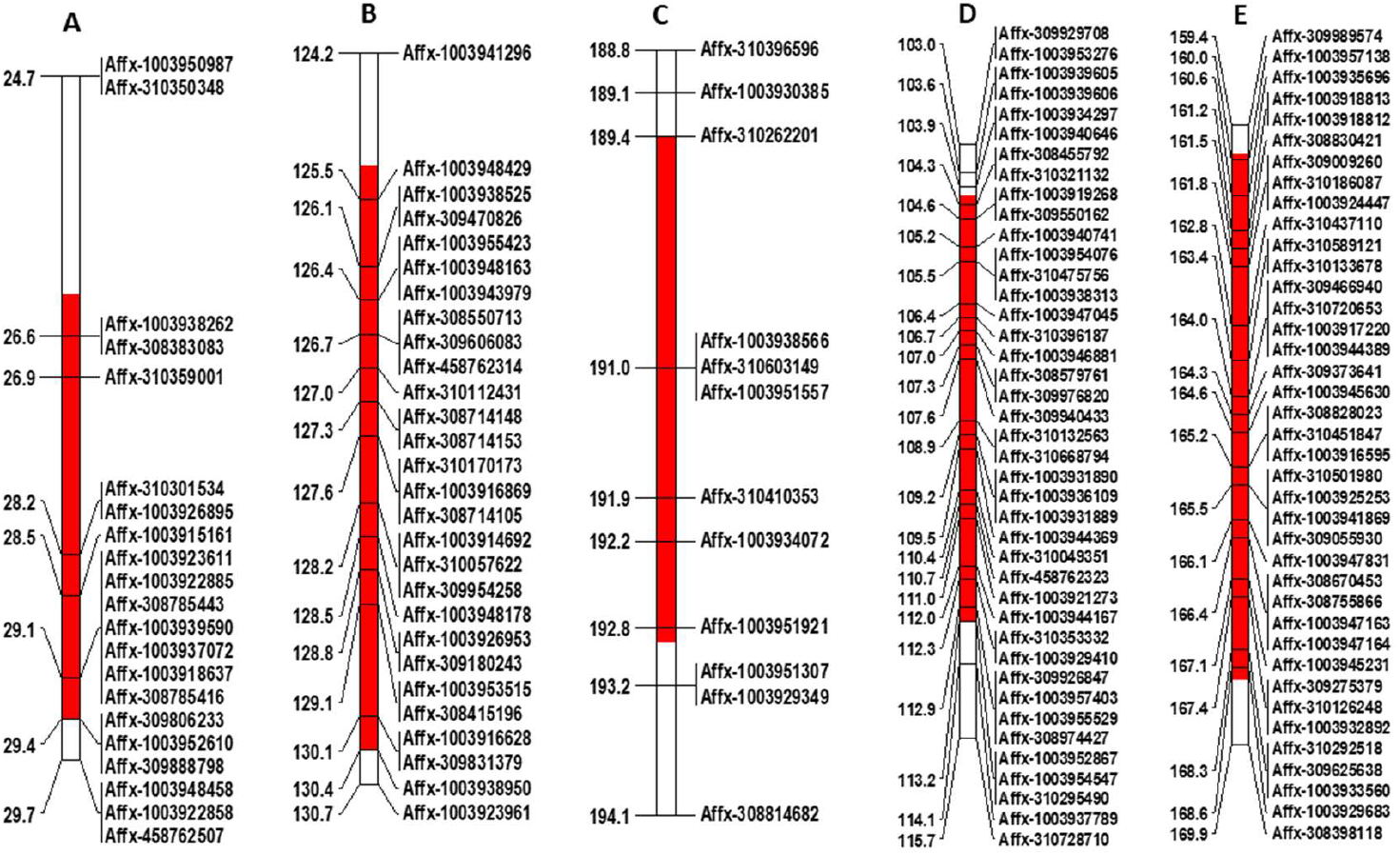
Location of chocolate spot resistance QTLs on chromosome 1 (A, B and C) and 6 (D and E) using 165 RILs derived from cross Mélodie/2 × ILB 938/2. The QTL positions are shown with a red bar, and the loci within the QTL regions are shown on the right of the bar. Only portions of the linkage map related to the QTL positions are displayed

### Identification of putative candidate genes

Analysis was conducted of the sequences of the SNP markers in the region of the two stable resistance loci, qBF1.1 and qBF1.2, which were identified by both QTL mapping approaches. A similarity search using BLASTn in the *M. truncatula* genome found a number of potential genes similar to the markers linked to the observed resistance against *B. fabae*. The sequence of SNP markers linked to qBF1.1 showed similarity with seven genes on chromosome 2 and those linked to qBF1.2 with 16 genes on chromosome 5 of *M. truncatula*, all of which were annotated as genes possibly associated with plant disease resistance and/or defence (Table 3).

**Table 3.**
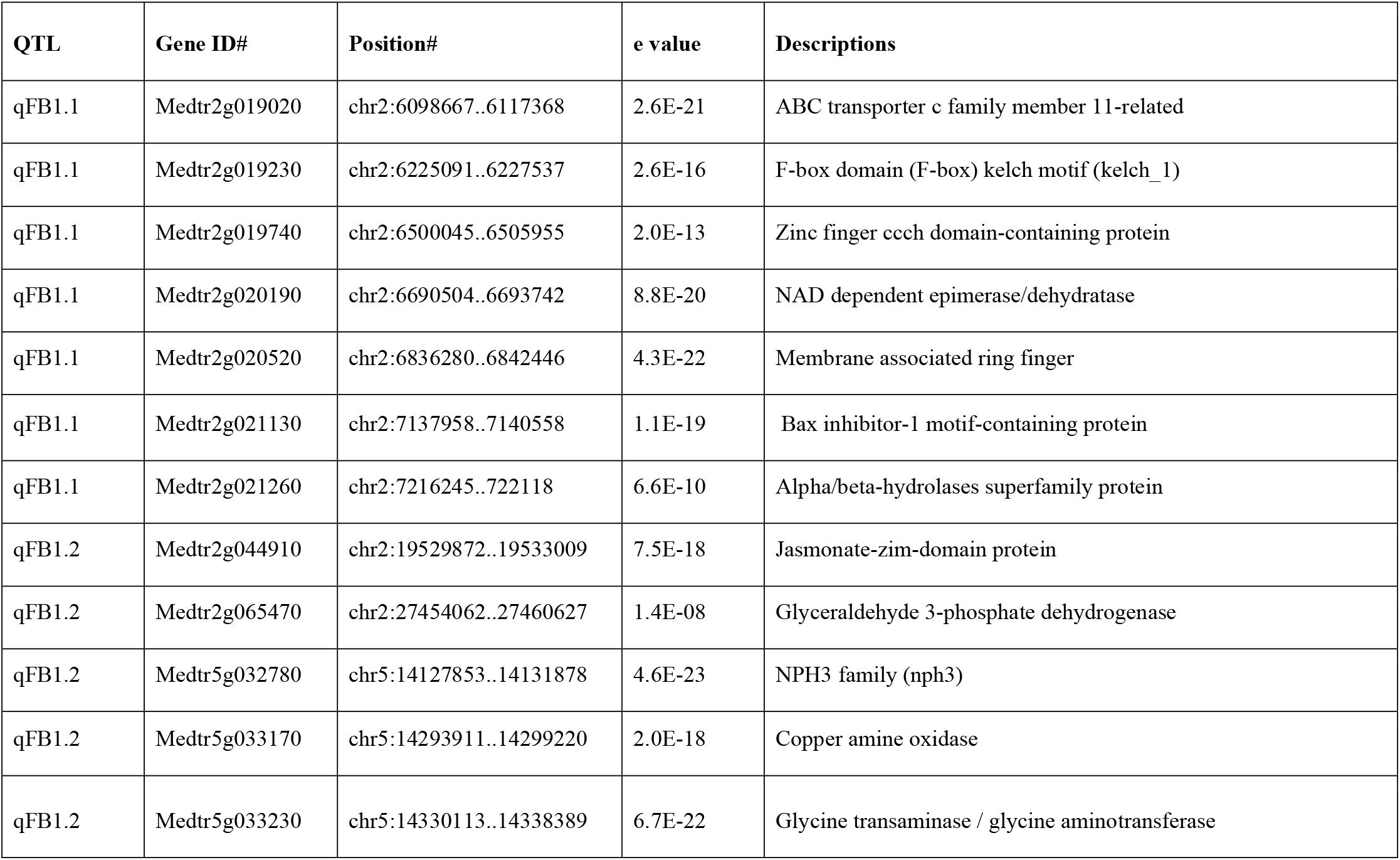
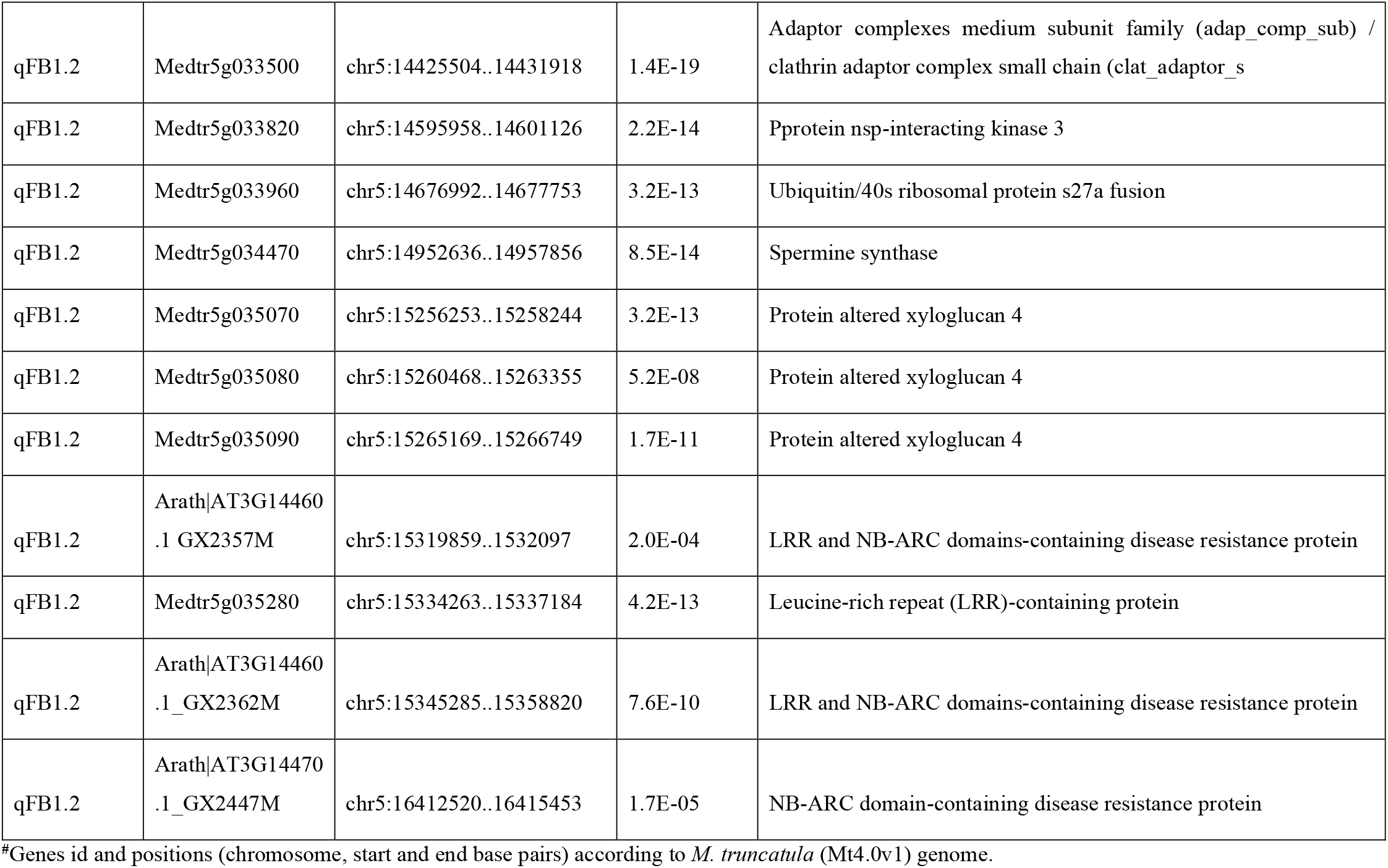
Candidate resistance and/or defense-related genes associated with QTLs for chocolate spot resistance based on BLASTn sequence similarity searches in the *Medicago truncatula* (Mt4.0v1) genome. The genes were queried using sequences of the SNP markers identified in the interval between the two stable resistance loci (qBF1.1 and qBF1.2).

| QTL | Gene ID# | Position# | e value | Descriptions |
| --- | --- | --- | --- | --- |
| qFB1.1 | Medtr2g019020 | chr2:6098667..6117368 | 2.6E-21 | ABC transporter c family member 11-related |
| qFB1.1 | Medtr2g019230 | chr2:6225091..6227537 | 2.6E-16 | F-box domain (F-box) kelch motif (kelch_1) |
| qFB1.1 | Medtr2g019740 | chr2:6500045..6505955 | 2.0E-13 | Zinc finger ccch domain-containing protein |
| qFB1.1 | Medtr2g020190 | chr2:6690504..6693742 | 8.8E-20 | NAD dependent epimerase/dehydratase |
| qFB1.1 | Medtr2g020520 | chr2:6836280..6842446 | 4.3E-22 | Membrane associated ring finger |
| qFB1.1 | Medtr2g021130 | chr2:7137958..7140558 | 1.1E-19 | Bax inhibitor-1 motif-containing protein |
| qFB1.1 | Medtr2g021260 | chr2:7216245..722118 | 6.6E-10 | Alpha/beta-hydrolases superfamily protein |
| qFB1.2 | Medtr2g044910 | chr2:19529872..19533009 | 7.5E-18 | Jasmonate-zim-domain protein |
| qFB1.2 | Medtr2g065470 | chr2:27454062..27460627 | 1.4E-08 | Glyceraldehyde 3-phosphate dehydrogenase |
| qFB1.2 | Medtr5g032780 | chr5:14127853..14131878 | 4.6E-23 | NPH3 family (nph3) |
| qFB1.2 | Medtr5g033170 | chr5:14293911..14299220 | 2.0E-18 | Copper amine oxidase |
| qFB1.2 | Medtr5g033230 | chr5:14330113..14338389 | 6.7E-22 | Glycine transaminase / glycine aminotransferase |
| qFB1.2 | Medtr5g033500 | chr5:14425504..14431918 | 1.4E-19 | Adaptor complexes medium subunit family (adap_comp_sub) / clathrin adaptor complex small chain (clat_adaptor_s |
| qFB1.2 | Medtr5g033820 | chr5:14595958..14601126 | 2.2E-14 | Pprotein nsp-interacting kinase 3 |
| qFB1.2 | Medtr5g033960 | chr5:14676992..14677753 | 3.2E-13 | Ubiquitin/40s ribosomal protein s27a fusion |
| qFB1.2 | Medtr5g034470 | chr5:14952636..14957856 | 8.5E-14 | Spermine synthase |
| qFB1.2 | Medtr5g035070 | chr5:15256253..15258244 | 3.2E-13 | Protein altered xyloglucan 4 |
| qFB1.2 | Medtr5g035080 | chr5:15260468..15263355 | 5.2E-08 | Protein altered xyloglucan 4 |
| qFB1.2 | Medtr5g035090 | chr5:15265169..15266749 | 1.7E-11 | Protein altered xyloglucan 4 |
| qFB1.2 | Arath\|AT3G14460<br>.1 GX2357M | chr5:15319859..1532097 | 2.0E-04 | LRR and NB-ARC domains-containing disease resistance protein |
| qFB1.2 | Medtr5g035280 | chr5:15334263..15337184 | 4.2E-13 | Leucine-rich repeat (LRR)-containing protein |
| qFB1.2 | Arath\|AT3G14460<br>.1 GX2362M | chr5:15345285..15358820 | 7.6E-10 | LRR and NB-ARC domains-containing disease resistance protein |
| qFB1.2 | Arath\|AT3G14470<br>.1 GX2447M | chr5:16412520..16415453 | 1.7E-05 | NB-ARC domain-containing disease resistance protein |

#Genes id and positions (chromosome, start and end base pairs) according to *M. truncatula* (Mt4.0v1) genome.

## Discussion

Resistance breeding is the most cost-effective and environmentally friendly strategy for controlling CS. Many studies have reported ILB 938 as a source of resistance to CS (e.g. Hanounik and Robertson 1988; Rhaïem et al. 2002; Villegas-Fernández et al. 2009; Maalouf et al. 2016). The introduction of CS resistance from this source into elite cultivars could be accelerated using MAS. To implement MAS in faba bean breeding programs, identification of markers linked to QTLs or genes conferring resistance to CS is the first step. In this study, the analysis of QTLs was conducted using a RIL population derived from ILB 938/2. The RIL population showed significant variation in pathogen response, as expected from the presence of resistance genes or alleles. The frequency distribution of the CS disease score in RILs showed a continuous distribution, indicating polygenic inheritance of CS resistance in ILB 938/2. To our knowledge, this study provides the first identification of genomic regions linked to CS resistance in this species.

Here we report the first linkage map, comprised of 4,089 SNP markers in six linkage groups that correspond to the six chromosomes of faba bean, generated for the ILB938-derived RIL population. The genetic linkage map spanned 1,229.5 cM with an average marker density of 0.3 cM. According to Stange et al. (2013), an increase in marker density from 5 to 1 cM could increase the power sufficiently to precisely localize and resolve closely linked QTLs. Therefore, the genetic map developed in this study has sufficient marker density to provide adequate power for QTL mapping. The total map size and the marker distributions along the LGs of the current genetic map are consistent with the consensus map previously published by Webb et al. (2016). It is shorter than the recently published SNP-based consensus map (1547.71 cM) by Carrillo-Perrillo et al. (2020). Like previous marker-based genetic maps (Webb et al. 2016; Carrillo-Perrillo et al. 2020) and cytogenetic studies (Lucretti et al. 1993; Doležel and Lucretti 1995), LG 1, corresponding to chromosome 1, contained the highest number of SNP markers and a larger size.

Using this genetic map and the CS phenotypic data for the RIL population, we mapped five QTLs associated with CS resistance on LG 1 and LG 6. Two QTLs on LG 1 (qBF1.1 and qBF1.2) with high LOD scores were detected by both WinQTLCart and R/qtl. These QTLs were derived from ILB 938/2 and explained a total phenotypic variation of 22%. The other three QTLs, one on LG 1 (qBF1.3) and two on LG 6 (qBF6.1 and qBF6.2), passed the threshold after permutation and were detected only by WinQTLCart. Interestingly, two of these three QTLs were inherited from the moderately susceptible parent Mélodie/2. In QTL studies for various traits in different crops, the inheritance of the resistance allele from a susceptible parent is not unusual, and a good example is the large effect QTL for improved yield in rice under drought conditions derived from the susceptible parent (Bernier et al. 2007).

When the sequences of the SNP markers identified in the regions of the two stable QTLs (qBF1.1 and qBF1.2) were used as BLASTn queries of the *M. truncatula* genome, the corresponding regions included several candidate genes that may play a role in disease resistance or defence against biotic stress. Of these, several are promising candidates: an alpha/beta-hydrolase protein, known as the Arabidopsis (*Arabidopsis thaliana*) Enhanced Disease Susceptibility 1 (EDS1) gene (Wittek et al. 2014; Vossa et al. 2019); Bax inhibitor-1 protein (Watanabe and Lam 2006); clathrin adaptor complex protein (Qiao et al. 2010); NAD-dependent epimerase/dehydratase (Islam et al. 2019); glyceraldehyde 3-phosphate dehydrogenase (Henry et al. 2015); NPH3 protein (Yang et al. 2016); spermine synthase (Janse van Rensburg et al. 2021); F-box protein (Hedtmann et al. 2017); and copper amine oxidase (Rea et al. 2002). All these genes are known to be involved in regulating programmed cell death against pathogens and in controlling cell death proliferation in response to reactive oxygen species (ROS). Recently, Castillejo et al. (2021) reported that resistance to *B. fabae* is linked to a more efficient photosystem II repair cycle in the resistant faba bean accession BPL 710. In response to *B. fabae* infection, different levels of ROS, lipid peroxidation, and activity of the enzymatic ROS scavenging system were detected in resistant and susceptible faba bean genotypes (El-Komy 2014). However, the antioxidative system is more effective in removing excess ROS produced during infection in the resistant genotype, and pathogenesis-related protein gene transcripts are expressed earlier and higher (El-Komy 2014; Castillejo et al. 2021). Genes encoding leucine-rich repeat-containing proteins such as NBS-LRR are typically disease-resistance (*R*) genes that detect the pathogen and activate downstream signaling, leading to pathogen resistance (Dodds and Rathjen 2010). We also identified jasmonate ZIM-domain proteins, which are involved in the responses to plant pathogens and abiotic stresses by regulating jasmonic acid signaling pathways and the cross-talk between various phytohormones (Zhou et al. 2015). Furthermore, the expression of genes encoding xyloglucan induces resistance against necrotrophic (*B. cinerea*) in Arabidopsis (Claverie et al. 2018).

Despite the fact that CS is one of the most damaging faba bean diseases worldwide, progress remains slow in breeding for CS resistance. Among the challenges related to faba bean improvement, the restricted appearance of the disease only during the humid climate cropping season makes traditional field-based phenotypic selection unreliable, particularly in dry areas like the Western Canadian prairies and parts of Australia (Adhikari et al. 2021; Khazaei et al. 2021b). This might explain why there were no QTL studies of CS previously reported. Therefore, conventional faba bean breeding could be augmented with molecular marker technologies to develop CS-resistant cultivars that are suitable for fulfilling the growing global demand for faba bean generated by its high seed protein content and great ecological service in cropping systems. Our study provides the first QTL identification for CS resistance using high-resolution SNP markers. Some of the identified QTLs in this work can be used as potential targets for further studies, and the linked markers can enable the possibility of using MAS in faba bean CS resistance breeding after validation in appropriate germplasm.

## Acknowledgements

The authors gratefully acknowledge funding from the Danish Innovation Fund for funding the NORFAB project (Innovation Fund Denmark grant number 5158-00004B); ADF (Agriculture Development Fund – Government of Saskatchewan, Canada); the Western Grains Research Foundation, Canada; the Saskatchewan Pulse Growers, Canada; and the Academy of Finland (decision 298314, “Papugeno”). We also thank Dr. Sabine Banniza (Pulse pathology lab, University of Saskatchewan) for technical advice and the staff of the Pulse Crop Breeding Programs for technical support at the University of Saskatchewan.

## Author contribution statement

HK, AV, FLS and AHS conceived and designed the experiments. TSG and MB performed the experiments. WC performed DNA extraction and library preparation. TSG and HK analyzed the data. AV, HK, FLS and AHS provided resources and acquired the funding. TSG wrote the first draft of the manuscript. All authors read and reviewed the manuscript.

## Figure captions

**Figure S1.**
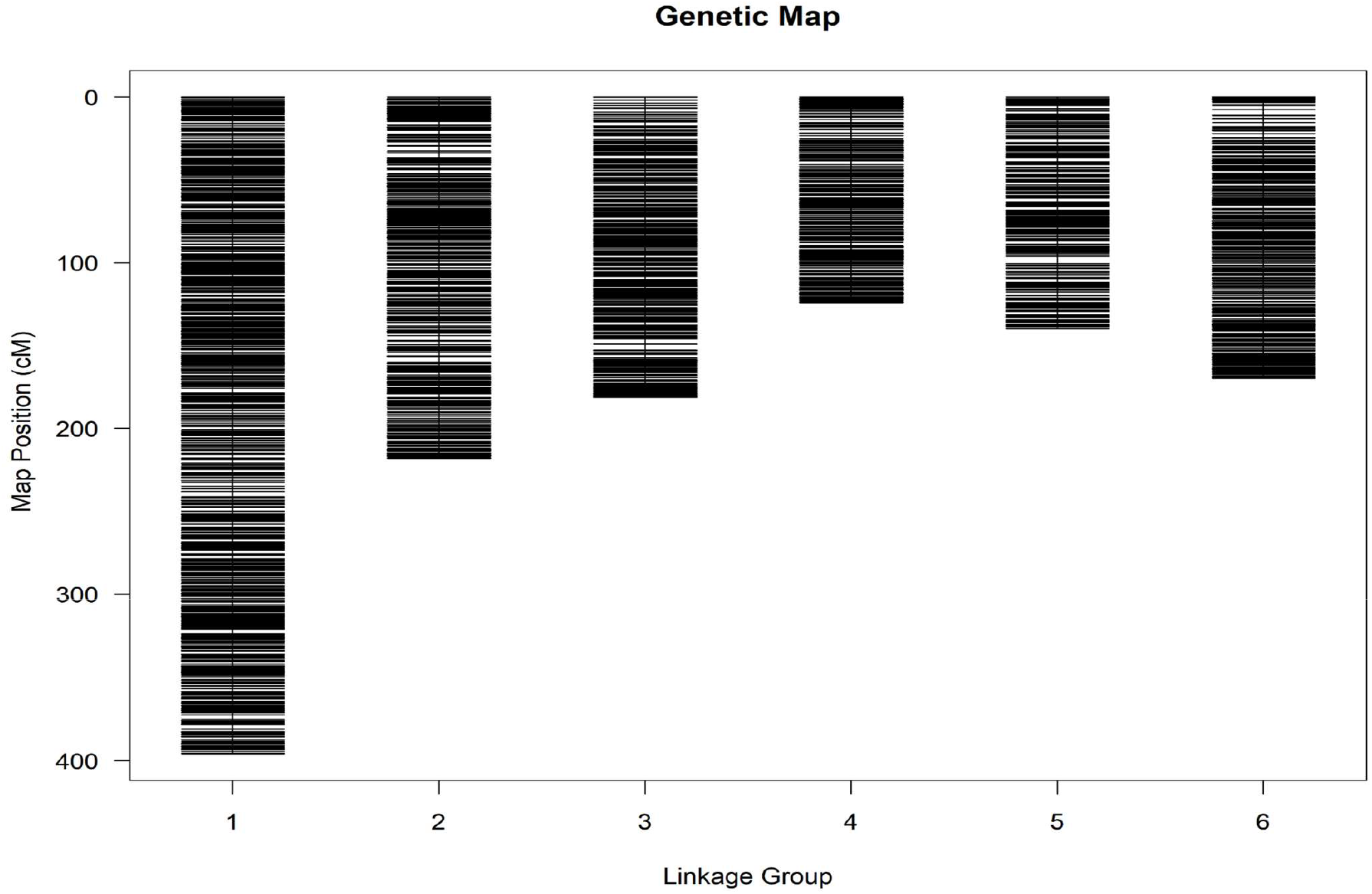
Genetic linkage map of cross Mélodie/2 × ILB 938/2 at F8.

